# *Manduca sexta* experience high parasitoid pressures in the field but minor fitness costs of consuming plant secondary compounds

**DOI:** 10.1101/2021.01.13.426617

**Authors:** Deidra J. Jacobsen

## Abstract

1. Plant-herbivore co-evolutionary interactions have led to a range of plant defenses that minimize insect damage and a suite of counter-adaptations that allow herbivores to feed on defended plants. Consuming plant secondary compounds results in herbivore growth and developmental costs but can have beneficial effects such as deterrence or harm of parasitoid enemies. Therefore, the role of secondary compounds on herbivore fitness must be considered in the context of the abundance and level of harm from natural enemies and the costs herbivores incur feeding on plant secondary compounds.
2. In this study, I combined field measurements of *Cotesia congregata* wasp parasitism pressure with detailed measurements of the costs of plant secondary compounds across developmental stages in the herbivore host, *Manduca sexta*.
3. I show that *C. congregata* parasitoids exert large negative selective pressures, killing 31-57% of *M. sexta* larvae in the field. *Manduca sexta* developed fastest during instars most at risk for parasitoid oviposition but growth was slowed by consumption of plant secondary compounds. The negative effects of consuming plant secondary compounds as larvae influenced adult size traits but there were no immune, survival, or fecundity costs.
4. These results suggest that developmental costs experienced by *M. sexta* herbivores consuming defensive compounds are minor in comparison to the strong negative survival pressures from abundant parasitoid enemies.

## Introduction

Coevolution between herbivorous insects and their host plants often mitigates their reciprocal (negative) fitness effects, resulting in rapid evolution of plant anti-herbivore defense and insect counter-adaptations to these defenses (Ehrlich and Raven 1964, Maron *et al*. 2019). Plants produce combinations of physical and chemical defenses that may lower herbivore fitness by reducing growth, disrupting development, decreasing survival, and/or attracting natural enemies of herbivores (Price *et al*. 1980, Howe and Jander 2008, Furstenberg-Hagg *et al*. 2013). Herbivores feeding on defended plants may experience immediate or delayed effects of consuming secondary compounds and these consequences may extend past the life stage at which the stress was experienced (Fellous and Lazarro 2010, van Dam *et al*. 2011). Understanding how insect herbivores evolve to feed on defended plants requires quantifying the fitness effects of plant defenses across herbivore life stages and examining the conditions under which consumption of plant secondary compounds is beneficial to herbivores. An important component of estimating these fitness consequences is to determine how additional stressors, such as natural enemies, affect herbivore fitness in conjunction with plant defense.

One way that herbivores mitigate the negative consequences of plant defense is if consumption of these secondary compounds directly or indirectly harms natural enemies. Insects are predicted to consume defended plants despite the apparent negative effects if the anti-enemy benefit of consuming plant secondary compounds outweighs the costs. Enemies can exert a large negative selective pressure on their targets when the enemies are at high abundance and/or when they drastically decrease herbivore fitness (Hassell and Waage 1984). Parasitoids, for instance, have an especially negative fitness effect because they kill their host at an immature (pre-reproductive) stage and reduce host fitness to zero (Godfray 1994). Endoparasitoids—those that develop inside the body of their hosts--are common enemies of lepidopteran species and endoparasitoid fitness may be particularly affected by host quality (Eggleton and Belshaw 1992). In insect hosts that sequester secondary compounds, there is clear co-option of plant toxins for herbivore anti-enemy defense but sequestration is considered more effective against predators rather than parasitoids because of the trade-off between chemical sequestration and immune function needed to defend against parasitoid eggs (Gauld *et al*. 1992, Smilanich *et al*. 2009).

In herbivore hosts that do not sequester plant secondary compounds, such as *Manduca sexta*, plant defenses can still reduce endoparasitoid success on hosts fed secondary compounds, either through direct toxicity to parasitoids or indirect effects on host quality (Beckage and Riddiford 1978, Thorpe and Barbosa 1986, Barbosa *et al*. 1991; Harvey *et al*. 2007). Endoparasitoids may come into contact with the toxic compounds their herbivore hosts consume as these compounds are detoxified or excreted from the host (Wink and Theile 2002, Kumar *et al*. 2014). Endoparasitoids are also sensitive to the indirect effects of secondary compounds on host growth, survival, and immune function (Parr and Thurston 1972, Price *et al*. 1980, Barbosa *et al*. 1991, Appel and Martin 1992, Stamp 1993, Alleyne and Beckage 1997, Ode 2006, Bukovinsky *et al*. 2009, Thaler *et al*. 2012, D’Incao *et al*. 2012). One host immune response in particular, the encapsulation and melanization of parasitoid eggs, can decrease parasitoid egg hatching success but whether this response is increased or hindered by secondary compounds is hard to predict given current evidence (Kraaijeveld *et al*. 2001, Bukovinszky *et al*. 2009, Smilanich *et al*. 2009).

Disentangling the effects of secondary compounds and their impacts on herbivore immune function and growth is necessary to determine how secondary compounds alter herbivore health and interactions between herbivores and parasitoids. Secondary compounds may prime the insect’s immune response to allow the insect to better respond to subsequent stress, such as parasitoid attack, or (alternatively) insect immunity may suffer as a result of eating defended plant tissues (Fellous and Lazzaro 2010). The impact of secondary compounds on herbivore immune function must be interpreted in the context of larval growth since limited resources are predicted to result in a trade-off between growth and immunity (Ode 2006, Bascunan-Garcia *et al*. 2010; van der Most *et al*. 2011, Wilson *et al*. 2019). The effects of secondary compounds may also be more pronounced at specific herbivore developmental stages. Studies that measure overall increases in time from hatching to pupation or short-term decreases in growth rate do not fully capture whether these developmental changes alter herbivore fitness or exposure to parasitoids (e.g. Parr and Thurston 1972, Granzow *et al*. 1985, Barbosa *et al*. 1991, Harvey *et al*. 2007, but see Van Dam *et al*. 2001).

Whether larval consumption of secondary compounds influences herbivore fitness depends in part on if stress experienced at the larval stage impacts adult mating and reproductive traits in addition to survival to adulthood (Bessin and Reagan 1990, Spurgeon *et al*. 1995). The effect of larval experience on reproductive fitness may differ based on insect life histories and developmental patterns. Larval experience has been shown to impact adult traits in non-holometabolous insects (Hopkins 1917, Corbet 1985), but has received less attention in insects that undergo metamorphosis because pupation is often thought of as a re-setting period that can erase larval experiences (Barron 2001, Fellous and Lazzaro 2010). However, pupal size and adult fecundity are correlated in some lepidopteran species, indicating that larval are not always negated by the restructuring that occurs during pupation and larval resource acquisition can affect adult traits (Bessin and Reagan 1990, Spurgeon *et al*. 1995, Kariyat and Portman 2016). Therefore, quantifying the impacts of larval stress on adult morphological and behavioral traits is necessary to determine if there are lasting effects of plant defense on adult mating and fitness traits or if larval consumption of secondary compounds simply alters survival to adulthood but the surviving adults are unaffected by larval stress.

In this study, I use the herbivore *Manduca sexta* (Lepidoptera: Sphingidae) to determine the fitness costs posed by natural enemies and the costs of larval consumption of chemically defended plants on herbivore immunity, development, survival, and adult fitness traits. By collecting and monitoring *M. sexta* from a field population, I establish that *Cotesia congregata* parasitoids exert large negative survival costs on larval *M. sexta*. Because the anti-parasitoid benefits of consuming host plants high in secondary compounds could outweigh mild negative developmental effects on herbivores, I also quantified the fitness effects of plant secondary compounds in field-collected and lab colonies of *M. sexta*. I show that while two different types of secondary compounds (inducible nicotine and constitutive rutin) affect *M. sexta* larval and adult size and morphological traits, they do not have strong negative effects on survival to adulthood, immune responses to artificial parasitoids, or adult fecundity. Because nicotine is known to have protective effects against *C. congregata* parasitoids (Beckage and Riddiford 1978, Thorpe and Barbosa 1986, Barbosa *et al*. 1991, Harvey *et al*. 2007), these results suggest that the developmental costs experienced by *M. sexta* consuming defensive compounds may be minor in comparison to the harm from abundant enemy pressures.

## Materials and Methods

### Study system: *Manduca sexta* and *Cotesia congregata*

*Manduca sexta* are ecologically and economically important pollinators and herbivores of Solanaceous plants. While feeding on host plants, *M. sexta* larvae are targeted by natural enemies, including *Braconid* wasp and *Tachinid* fly parasitoids that lay eggs inside their hosts (Yamamoto and Fraenkel 1960, Stireman *et al*. 2006, Garvey *et al*. 2020). *Cotesia congregata* parasitoid eggs hatch and feed inside the *M. sexta* host larvae before they emerge from the host larval cuticle to pupate, ultimately killing the host (Alleyne and Beckage 1997). Prior studies have shown that consumption of plant secondary compounds by larvae of *M. sexta* can be protective against parasitoids by deterring parasitoid oviposition and harming parasitoid development (Beckage and Riddiford 1978, Thorpe and Barbosa 1986, Barbosa *et al*. 1991; Harvey *et al*. 2007).

### Field collection of *M. sexta* larvae

*Manduca sexta* larvae were collected from leaves of dark tobacco (*Nicotiana tabacum*) at the University of Kentucky Research and Education Center (Princeton, KY) to determine parasitoid abundance and establish a field-collected colony for experiments testing the effects of secondary compounds on *M. sexta*. The 4.5-acre field area contained ~4900 dark tobacco plants/acre, with a cured leaf content of approximately 3-5% nicotine (Dr. Andrew Bailey, personal communication). Over three field collection dates all larvae found in the field were collected, for a total of 395 *M. sexta* larvae ranging from second to fifth/sixth instar collected (21 July 2013 N = 98; 20 August 2013 N = 156; 28 July 2014 N = 141). These dates were timed to occur after the residual insecticide used in transplanting (late May-early June) wore off and before application of additional pesticides.

### Measurements of parasitoid abundance on field *M. sexta*

After each of the three field collections, *M. sexta* larvae were brought back to the lab and monitored twice daily for parasitoid emergence. Because *M. sexta* consume a large amount of leaf tissue, field-collected larvae were transitioned to an artificial wheat germ-based diet with 10-20% wet volume of Solanaceous leaf tissue added to facilitate diet acceptance (SI Table 1). Larvae were fed *ad libitum* under 14:10 light:day cycles at 22.2+/-0.5 °C (Bell *et al*. 1975). Parasitoid development takes a predictable number of days, meaning that the time between field collection and parasitoid emergence can be used to estimate the instar at which parasitoid oviposition occurred (Gilmore 1938). In July 2014, I recorded the approximate instar at field collection and determined the time it took parasitoids to emergence from different host instars.

Chi-squared tests were used to test for variation in the proportion of *M. sexta* larvae with parasitoids among the three field collection dates. Parasitoid load and number of days post-field collection until *C. congregata* emergence were compared for larvae collected from the field at different instars using Poisson general linear models (GLM) (R v. 3.2.2; R Core Team 2014). Robust standard errors were used as Breusch-Pagan tests showed heteroskedasticity (bptest() in LMTEST; Zeileis and Hothorn 2002). Instar five was excluded from instar-specific models because I collected only two fifth instars.

### Rearing of *Manduca sexta* field-collected and lab colonies

Because laboratory and natural populations of *M. sexta* have been shown to have different evolutionary histories and responses to stressful conditions (Kingsolver 2007, Diamond *et al*. 2010, Kingsolver *et al*. 2020), I used the surviving *M. sexta* from the 2014 field collection to establish a field-collected colony to use alongside the standard lab colony to test the effects of secondary compounds on herbivore growth and fitness (Supplementary Methods 1). The lab colony was derived from a colony maintained under solely laboratory conditions (artificial diet, no introduction of wild individuals) for >250 generations since the 1960s (Carolina Biological Supply, Kingsolver *et al*. 2009). Field-collected and lab colonies were kept in separate cages in the same greenhouse. Prior to experiments, the field-collected colony was reared through a generation on solely artificial diet to control for maternal effects.

### Preparation of *M. sexta* diets with secondary compounds

Using artificial diets containing either nicotine or rutin (SI Table 1), I tested the effects of secondary compounds on *M. sexta* development and fitness traits. As a specialist herbivore, *M. sexta* often feed on leaves containing nicotine, a pyridine alkaloid found only in the Solanaceae plant family, which serves as a defensive chemical against herbivory and can be induced via the jasmonic acid pathway (Keinanen *et al*. 2001, Steppuhn *et al*. 2004). I used 0.5% wet weight nicotine, which represents a high but relevant concentration that larvae feeding on tobacco would encounter (Parr and Thurston 1972, Saitoh *et al*. 1985, Sisson and Saunders 1982, Sisson and Saunders 1983, Thompson and Redak 2007). To test whether herbivore responses to plant defensive chemicals are consistent across different compounds, I also tested the effects of 0.5% rutin (quercetin 3-rhamnoglucoside) on the same herbivore traits. Rutin is found in 32 plant families and is constitutively present at 0.008-0.61% wet mass in tobacco (Krewson and Naghski 1953, Keinanen *et al*. 2001, Kessler and Baldwin 2004). *Manduca sexta* used for the immunity, growth, and adult measurements were fed artificial diet (control, 0.5% nicotine, or 0.5% rutin depending on treatment) *ad libitum*.

### *M. sexta* larval immune responses to artificial parasitoids

Injections of artificial parasitoid eggs into *M. sexta* larvae were used to test whether secondary compounds alter host immune responses and if growth and immunity trade off. *Manduca sexta* from both colonies were collected concurrently as neonate larvae and reared individually on 0.5% nicotine, 0.5% rutin, or control diets until the fourth instar (N = 27-30 per diet treatment for the lab colony and N = 16-20 per diet treatment for the field-collected colony). Fourth-instar larvae were used for injections because larvae are large enough to manipulate without causing death (Beetz *et al*. 2008). Forceps were used to insert an artificial parasitoid egg (a 2 mm-long piece of roughened nylon filament) through a needle hole pricked behind the fourth proleg as in Piesk *et al*. (2013). Larvae were returned to their respective diets and fed readily after egg insertion. Pre-challenge growth rate was calculated as ln(larval mass at fourth instar)/number of days from hatching to fourth instar. Post-challenge growth rate was calculated as ln(larval mass 24 hours post egg insertion/larval mass at time of egg insertion) (Diamond and Kingsolver 2011).

After the final weighing, larvae were frozen at −20°C for dissections to quantify the strength of the immune response to the artificial parasitoid. Melanization (dark buildup by hemocyte immune cells) was photographed using a Leica M205FA Stereo microscope and the percent melanized was calculated using ImageJ (Diamond and Kingsolver 2011). Percent melanized was used for GLM with robust standard errors to determine whether immune responses differed based on secondary compounds, prior condition (pre-challenge growth rate), or trade-offs between post-challenge growth rate and melanization. Pairwise interactions between diet-colony and diet-growth rates were non-significant (*P* > 0.1 for all) and were removed from the model. Area melanized was transformed for non-normality using Box-Cox lambda power transformations after scaling of non-positive values (BC = 0.1; boxcox() in MASS; Box and Cox 1964).

### *M. sexta* larval and pupal traits on nicotine and rutin diets

Because of the large numbers of *M. sexta* needed, the effects of nicotine and rutin on growth and fitness traits were tested and analyzed at separate generations. Larvae from both colonies were fed control or experimental diets (0.5% nicotine or 0.5% rutin) and monitored daily for molting and the number of days per larval instar (N = 80 per diet treatment and colony). The total number of larval instars was recorded because larvae undergo either five or six instars depending on size (Kingsolver 2007) (SI Table 2). Poisson GLM and Wald tests were used to test for variability in the number of days per instar and whether any instar was more sensitive to the effects of nicotine or rutin (wald.test() in AOD; Lesnoff and Lancelot, 2012). Non-significant interactions (*P* > 0.1) between instar and nicotine or rutin were removed.

I recorded the number of days to pupation and pupal mass for *M. sexta* males and females to test whether larval consumption of nicotine or rutin altered pupal traits. Poisson GLM was used to determine if nicotine or rutin extended the number of days to reach pupation. ANOVA with type III sums-of-squares was used to test for differences in pupal size mass based on secondary compounds or sex and whether the effects of nicotine or rutin were stronger for either sex (diet*sex interaction) (Anova() in CAR; Fox and Weisberg 2011). Pupal mass for lab moths in the rutin experiment was transformed by BC = 2. One-sided Fisher tests (fisher.test()) were used to test if secondary compounds increased larval and pupal deformities.

### *M. sexta* adult size and fitness traits

To test if larval consumption of secondary compounds resulted in size differences post-pupation, surviving adults were frozen at −20°C the morning post-eclosion to measure adult body and wing length. Kaplan-Meier survival analyses were used to determine if either secondary compound reduced moth survival to eclosion (survdiff() in SURVIVAL; Therneau and Grambsch 2000, Therneau 2015). ANOVA models for adult body and wing length included diet, sex, and a diet* sex interaction. Wing length was transformed by BC = 6 for moths in the lab colony.

Fecundity estimates were obtained by dissecting ovarioles from adult females and counting follicle numbers under a dissecting scope as in Diamond *et al*. 2010 (N = 25-30 per diet treatment for the lab colony and N = 8-25 per diet treatment for the field-collected colony). Because larger moths may produce more eggs, the ratio of follicles to body area was used as the dependent variable in ANOVA models testing for an effect of larval consumption of secondary compounds on fecundity. Follicles/body area was calculated as: (number of follicles in a female moth) / (½ body length x ½ body width x 3.14). Correlations among adult traits are shown in SI Table 3.

### *M. sexta* larval dietary choice trials

Binary choice trials were used to determine if neonate *M. sexta* exhibit a preference for artificial diets with or without secondary compounds. Neonate larvae from both colonies were collected within three hours of hatching and placed in the center of individual 9 cm diameter petri dishes with 1 cm^2^ pieces of control diet and experimental diet (0.5% nicotine or 0.5% rutin) placed on opposite sides. Dishes were oriented haphazardly under 14:10 dark:light conditions and monitored at 1, 6, and 24 hours before scoring contact with either diet at 48 hours as a choice. Larvae did not leave or switch once choosing a diet. Chi-Squared analyses were used to test if control or experimental diets were chosen significantly more than half the time. Larvae that did not choose in 48 hours (field N = 11/60 and lab N = 3/76) were excluded.

## Results

### *Cotesia congregata* parasitoids are common on *M. sexta* larvae in the field

Surveys of a tobacco plot at three timepoints revealed that a high but variable proportion of *M. sexta* larvae were parasitized. The highest proportion of larvae parasitized by *C. congregata* was observed at the first collection (0.57 parasitized in July 2013), compared with 0.31 parasitized in August 2013 and 0.39 parasitized in July 2014 (*X*^2^_2_ = 16.74, *P* < 0.001) (SI Fig. 1). Median *C. congregata* parasitoid load emerging from an individual larva was 21.5 (N = 44) and all larvae with parasitoids emerging died before pupation. Tachinid fly parasitoids eclosed from only two *M. sexta*.

The timing of *C. congregata* parasitoid emergence from *M. sexta* collected in the field at different instars was consistent with parasitoids ovipositing in younger larvae. Because parasitoids take a predictable amount of time to emerge from the host cuticle after oviposition (Gilmore 1938), the length of time between field collection and parasitoid emergence for larvae of different instar stages can be used to determine the age at parasitism. The number of days post-*M. sexta* field collection to *C. congregata* emergence was higher for host *M. sexta* collected as younger instars (GLM; instar 2 *b* = 2.599, *P* < 0.01; instar 3 *b* = −0.629, *P* < 0.001; instar 4 *b* = −1.100, *P* < 0.001). There were no significant increases in the number of parasitoids emerging from third and fourth instar host *M. sexta* larvae compared with second instar host larvae (GLM; instar 2 *b* = 2.99, *P* < 0.01; instar 3 *b*= 0.255, *P* = 0.236; instar 4 *b* = 0.340, *P* = 0.094) (Table 1).

**Table 1.** Proportion of *Manduca sexta* with parasitoids, parasitoid load, and emergence times on *M. sexta* collected from the field as different instars. The proportion of larvae with parasitoids was calculated based on field collections from July 2014. The number of *C. congregata* is presented as the median per *M. sexta* host. The time to emergence is presented as the median number of days from *M. sexta* field collection to parasitoid larval emergence through the host cuticle. For *C. congregata* number and days to emergence, host sample size and inter-quartile range are presented in parentheses.

| *M. sexta* instar | Proportion of larvae with parasitoids | Median number of *C. congregata* per host | Days until *C. congregata* emergence |
| --- | --- | --- | --- |
| **2** | 0.20 (N=56) | 18 (N=11, IQR= 13-26) | 14 (N=11, IQR= 12-15) |
| **3** | 0.53 (N=53) | 20 (N=21, IQR= 12-30) | 7 (N=58, IQR= 6-9) |
| **4** | 0.60 (N=20) | 28 (N=12, IQR= 18-36) | 4 (N=23, IQR= 3-5) |

### Immune responses to an artificial parasitoid do not trade-off with larval growth

Following implantation of an artificial parasitoid egg, *Manduca sexta* immune response (melanization) was higher in larvae with faster growth rates, regardless of diet. There was a positive relationship between growth rate post-challenge and the melanization of the artificial parasitoid (GLM; z = 0.20, *P* < 0.001). Growth rate prior to the artificial parasitoid did not affect melanization (z = −0.03, *P* = 0.85). Neither nicotine (z = −0.02, *P* = 0.37) nor rutin (z = 0.04, *P* = 0.10) affected melanization. Larvae from the field-collected colony had higher immune responses to the artificial parasitoid than larvae from the lab colony (z = 0.080, *P* < 0.001) with a mean melanization level of 19% for the field-collected colony and 12% for the lab colony.

### Secondary compounds increase developmental time for each larval instar

Developmental assays of *M. sexta* larvae revealed that the number of days needed to complete each instar is variable and secondary compounds extend the length of each instar. Nicotine and rutin increased the number of days needed to complete each of the first four instars but specific instars were not more sensitive to the effects of the secondary compounds (Table 2). Larvae spent the fewest number of days in the second instar but there was variation in development times between the nicotine and rutin experiments. In the *M. sexta* generation used to test the effects of nicotine, the number of days taken to complete the second and third instars was shorter than the number of days taken to complete the first and fourth instars (Table 2A). In the generation used to test the effects of rutin, only the second instar was shorter (Table 2B).

**Table 2.** The number of days spent in each instar in both the field-collected colony and lab colony increased in response to A) nicotine or B) rutin. Grey rows show the total number of days these secondary compounds add to the number of days per instar and significant effects of nicotine and rutin are indicated by asterisks (*). Rows for each instar indicate the total number of days spent in each instar and the results of the GLM and Wald tests comparing the lengths of instars 2,3, and 4 to the time spent in instar 1. Asterisks (*) following *P* values for instars show significantly shorter or longer times in those instars compared to instar 1 (*P* < 0.05).

| A) | <u>Field</u> |  |  | <u>Lab</u> |  |  |
| --- | --- | --- | --- | --- | --- | --- |
|  | days | *z* | *P* | days | *z* | *P* |
| **Nicotine** | +0.7/instar | 2.77 | <0.01* | +0.7/instar | 3.53 | <0.01* |
| instar 1 | 5.4 | 35.31 | <0.01 | 4.7 | 35.40 | <0.01 |
| instar 2 | 4.1 | -4.30 | <0.01* | 3.6 | -4.40 | <0.01* |
| instar 3 | 4.0 | -4.80 | <0.01* | 4.0 | -2.62 | 0.01* |
| instar 4 | 5.0 | -1.22 | 0.22 | 4.7 | 0.00 | 1 |
| B) | <u>Field</u> |  |  | <u>Lab</u> |  |  |
|  | days | *z* | *P* | days | *z* | *P* |
| **Rutin** | +0.8/instar | 2.57 | 0.01* | +0.7/instar | 3.68 | <0.001* |
| instar 1 | 4.6 | 23.45 | <0.01 | 4.2 | 32.80 | <0.01 |
| instar 2 | 3.9 | -3.06 | 0.01* | 3.5 | -2.95 | <0.01* |
| instar 3 | 4.1 | -1.24 | 0.22 | 4.3 | 0.50 | 0.62 |
| instar 4 | 5.7 | 2.68 | <0.01* | 5.3 | 4.11 | <0.001* |

The overall effect of secondary compounds on larval development was to increase the number of days from hatching to pupation. In both colonies, the number of days from hatching to pupation was higher on the nicotine diet (GLM; field nicotine z = 2.183, *P* = 0.029; lab nicotine z = 4.039, *P* < 0.001) and on the rutin diet (field rutin z = 4.149, *P* < 0.001; lab rutin z = 5.65, *P* < 0.001) compared to control diets (Table 3). Larvae normally complete five instars, but a small percent of larvae went through an additional sixth instar prior to pupation and this was more common in *M. sexta* in the lab colony fed nicotine and for *M. sexta* in lab and field-collected colonies fed rutin (SI Table 2).

**Table 3.**
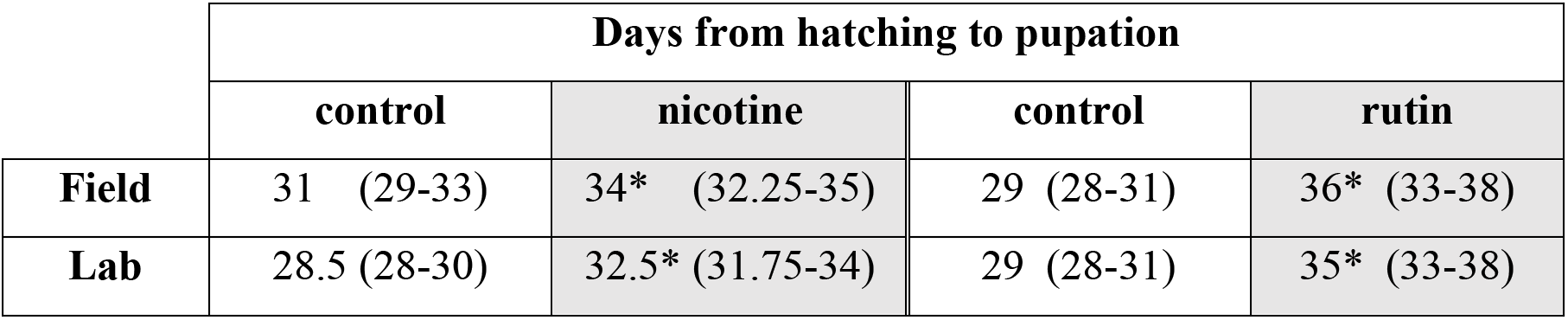
Consumption of nicotine and rutin increased the number of days to pupation for both the field-collected and lab colonies. Values are presented as medians followed by interquartile ranges in parentheses. Asterisks (*) indicate a significantly longer development time on the diet containing the secondary compound compared with the control (GLM *P* < 0.05). Separate controls are presented for nicotine and rutin because the effects of these secondary compounds were tested at different generations.

### Pupal mass was reduced by larval consumption of secondary compounds

Larval consumption of secondary compounds decreased pupal mass but the sex-specific patterns differed between nicotine and rutin (Fig. 1). Pupal mass was smaller when larvae were fed nicotine in both colonies (ANOVA; field nicotine *F*_1, 76_ = 21.562, *P* < 0.001; lab nicotine *F*_1, 112_ = 23.527, *P* < 0.001) (Fig. 1A). There was an interaction between sex and nicotine in the lab colony, such that females had a greater decrease in pupal mass from nicotine than males (lab sex*nicotine *F*_1,112_ = 5.285, *P* = 0.023) and male pupae were smaller than female (lab sex *F*_1,112_ = 8.265, *P* = 0.005). The field-collected colony had no differences between male and female pupal mass (field sex *F*_1, 7_6 = 0.479, *P* = 0.491) and there was no interaction between sex and nicotine (field sex*nicotine *F*_1, 76_ = 0.176, *P* = 0.676) (Fig. 1A). Pupal mass also decreased in both colonies in response to rutin (ANOVA; field rutin *F*_1, 45_ = 7.362, *P* = 0.009; lab rutin *F*_1, 118_ = 7.975, *P* = 0.006). There was no interaction between sex and rutin (field sex*rutin *F*_1,45_ = 2.196, *P* = 0.145; lab sex*rutin F_1, 118_ = 0.277, *P* = 0.600) although male pupae were smaller than female (field sex *F*_1, 45_ = 7.071, *P* = 0.011; lab sex *F*_1, 118_ = 8.024, *P* = 0.005) (Fig. 1B).

**Figure 1.**
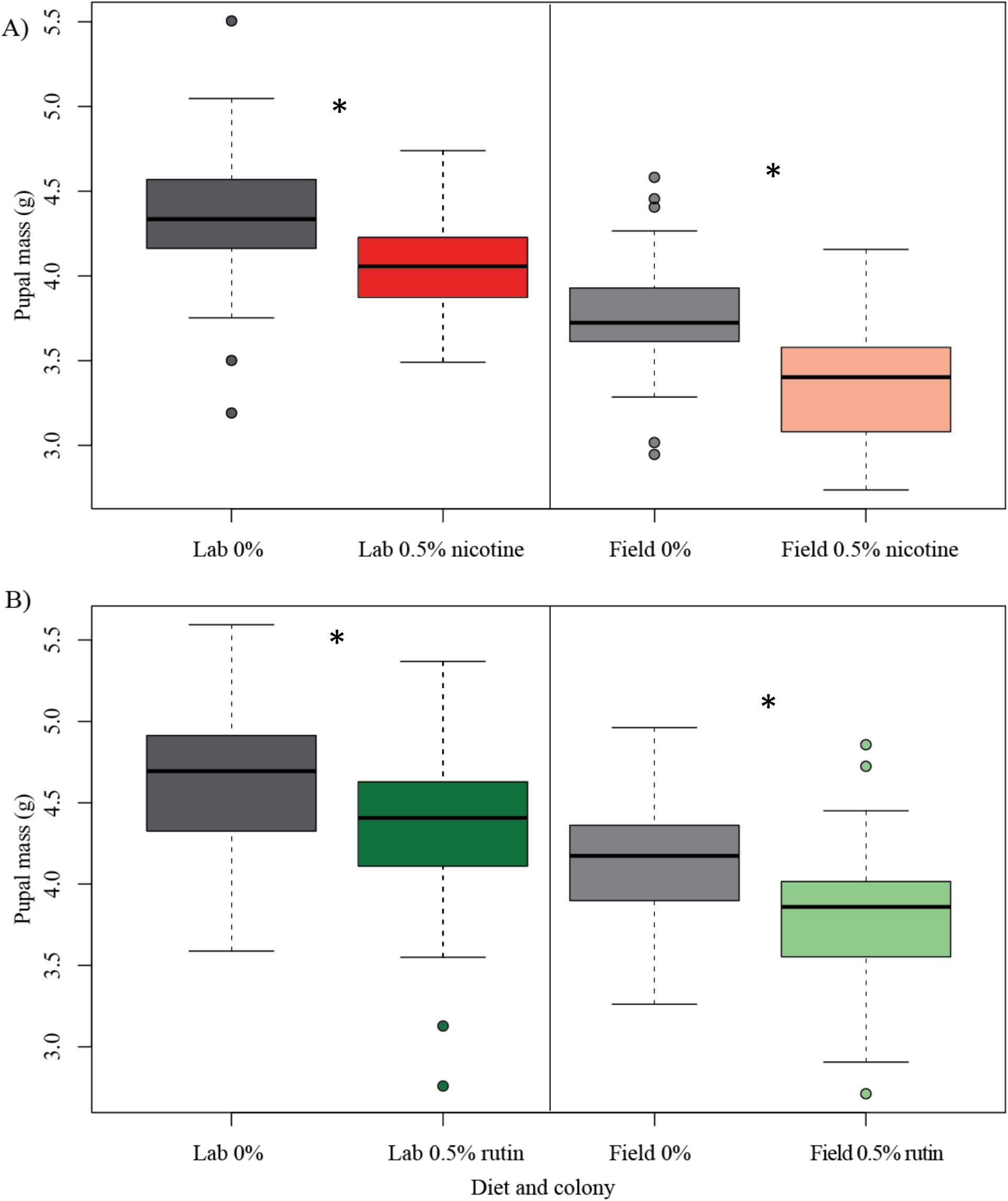
*Manduca sexta* pupae were smaller when larvae were fed secondary compounds. **A)** *M. sexta* larvae fed nicotine (red) weighed less at pupation than those fed control diets (grey) (ANOVA, *P* < 0.01 for both colonies) **B)** *M. sexta* larvae fed rutin (green) weighed less at pupation than those fed control diets (grey) (ANOVA, *P* < 0.01 for both colonies). Separate controls are presented for nicotine and rutin because the effects of these secondary compounds were tested at different generations.

### Secondary compounds do not increase *M. sexta* deformities

Minor deformities at the larval and pupal stage are common during *M. sexta* development but were not increased by dietary nicotine or rutin. The main deformities observed were incomplete larval molting (field N = 8/350 larvae; lab N = 13/320 larvae) and incomplete sclerotization of pupal cases (field N = 1/131 larvae; lab N = 26/243 larvae). The incidence of deformities was not increased by nicotine (one-sided Fisher’s exact tests; molting: field *P* = 0.75, lab *P* = 0.5; incomplete sclerotization: field *P* = 1, lab *P* = 0.67) or by rutin (one-sided Fisher’s exact tests; molting: field *P* = 0.34, lab *P* = 0.36; incomplete sclerotization: field *P* = 0.48, lab *P* = 0.51).

### *M. sexta* survival to adulthood is not decreased by larval consumption of secondary compounds

The proportion of *M. sexta* surviving to adult eclosion was not significantly reduced by either secondary compound in the larval diets. Larvae from the lab colony had only marginally significantly reduced survival to adult eclosion when reared on nicotine compared to those fed the control diet (*X*^2^_1_ = 3.6, N = 149, *P* = 0.057) and there were no differences in survival for moths from the field-collected colony when fed nicotine versus control diets (*X*^2^_1_ = 0, N = 152, *P =* 0.886) (Fig. 2A). Rutin did not significantly decrease survival of moths from either colony (lab: *X*^2^_1_ = 0.7, N = 158, *P* = 0.408; field *X*^2^_1_ = 0, N = 189, *P* = 0.956) (Fig. 2B).

**Figure 2.**
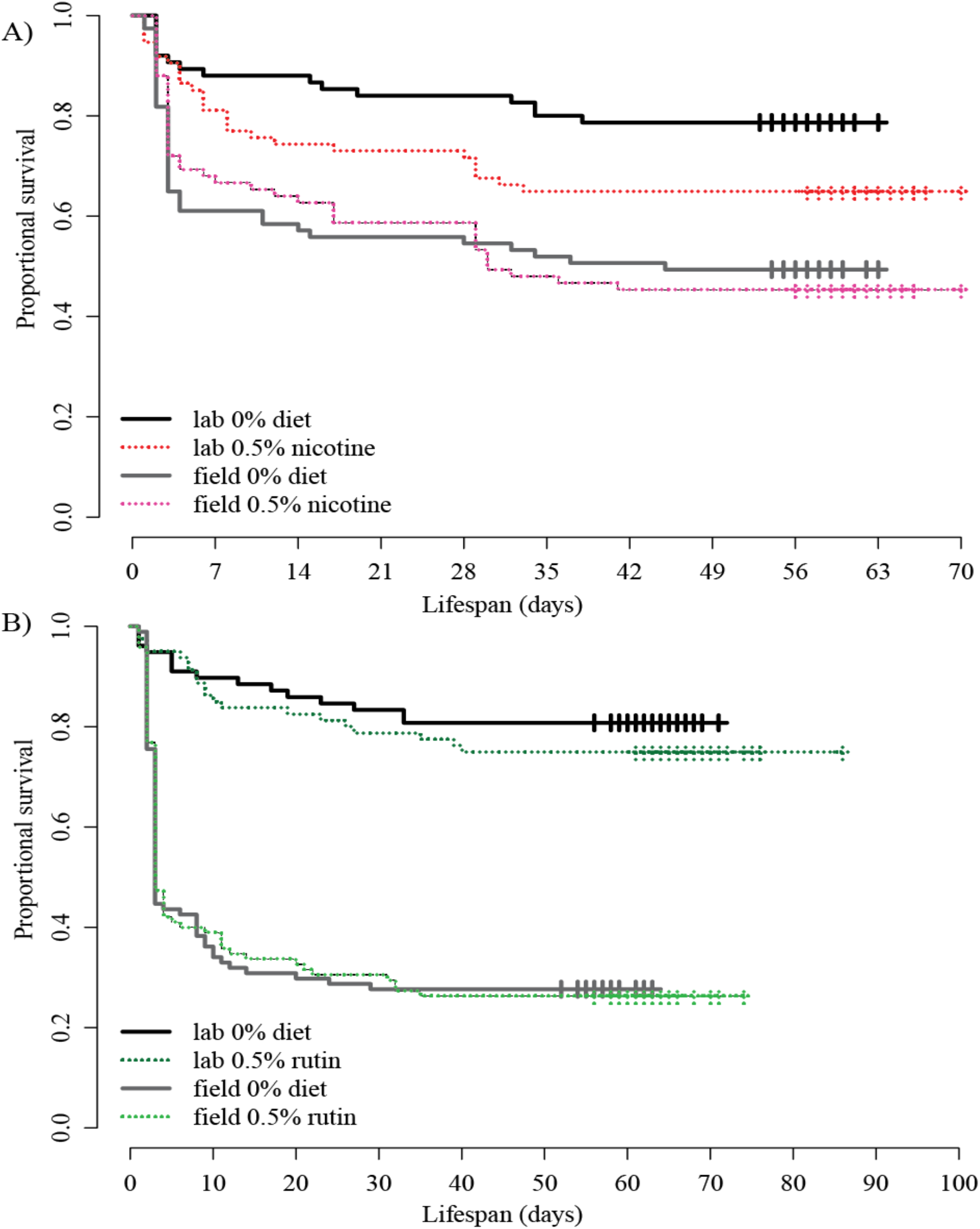
Secondary compounds did not significantly decrease *Manduca sexta* survival to adult eclosion. **A)** *M. sexta* from the field-collected colony had similar survival on nicotine (red) versus control diets (grey) but there was a slight, non-significant decrease in survival for lab colony fed nicotine (Kaplan-Meier survival curves; field *P* = 0.9, lab *P* = 0.06). **B)** Neither colony had reduced survival on rutin (green) versus control diets (grey) (field *P* > 0.01, lab *P* > 0.1). Tick marks represent censored data (insects that pupated prior to end of time period).

### Adult body size was smaller when moths had consumed secondary compounds as larvae

Measurements of body length on newly eclosed adults showed that larval consumption of secondary compounds decreased *M. sexta* size at the adult stage, but the effects differed for female and male moths. These effects are not explained by size variation between the sexes, as male and female adult body lengths were similar within a colony (nicotine experiment: field sex *F*_1, 72_ = 0.342, *P* = 0.561; lab sex *F*_1, 105_ = 0.398, *P* = 0.530; rutin experiment: field sex F_1,43_ = 1.003, *P* = 0.322; lab sex F_1,115_ = 01.617 *P* = 0.206).

For both colonies, nicotine decreased adult length (ANOVA: field nicotine *F*_1, 72_ = 7.978, *P* = 0.006; lab nicotine *F*_1, 105_ = 19.896, *P* < 0.001). The negative effect of nicotine on adult length was stronger on females than males from the field-collected colony (field sex*nicotine *F*_1, 72_ = 4.072, *P* = 0.047). There was no sex difference in the effect of nicotine in the lab colony (lab sex*nicotine *F*_1, 105_ = 0.366, *P* = 0.546).

Larval consumption of rutin decreased adult male body size in the field-collected colony (field sex*rutin F_1,43_ = 4.419, *P* = 0.041; field rutin F_1,43_=3.835, *P* = 0.057). There was no effect of rutin on adult moth size for the lab colony (lab sex*rutin F_1,115_ = 0.125, *P* = 0.724; lab rutin F_1,115_ = 2.878, *P* = 0.093).

### Adult wing size was smaller when moths had consumed secondary compounds as larvae

Larval consumption of secondary compounds decreased adult wing size. Males had smaller wings than females but were not more sensitive to the effect of secondary compounds on wing length.

For both colonies, wing length was smaller when the *M. sexta* had been fed nicotine as larvae (ANOVA; field nicotine *F*_1, 70_ = 19.432, P<0.001; lab nicotine *F*_1, 102_ = 18.505, *P*<0.001). Although males had smaller wings than females (field sex *F*_1, 70_ = 34.753, *P*<0.001; lab sex *F*_1, 102_ = 73.822, *P*<0.001), the effect of nicotine did not differ between the sexes within a colony (field sex*nicotine *F*_1, 70_ = 0.478, *P* = 0.492; lab sex*nicotine *F*_1, 102_ = 1.730, *P* = 0.191) (Fig. 3A).

**Figure 3.**
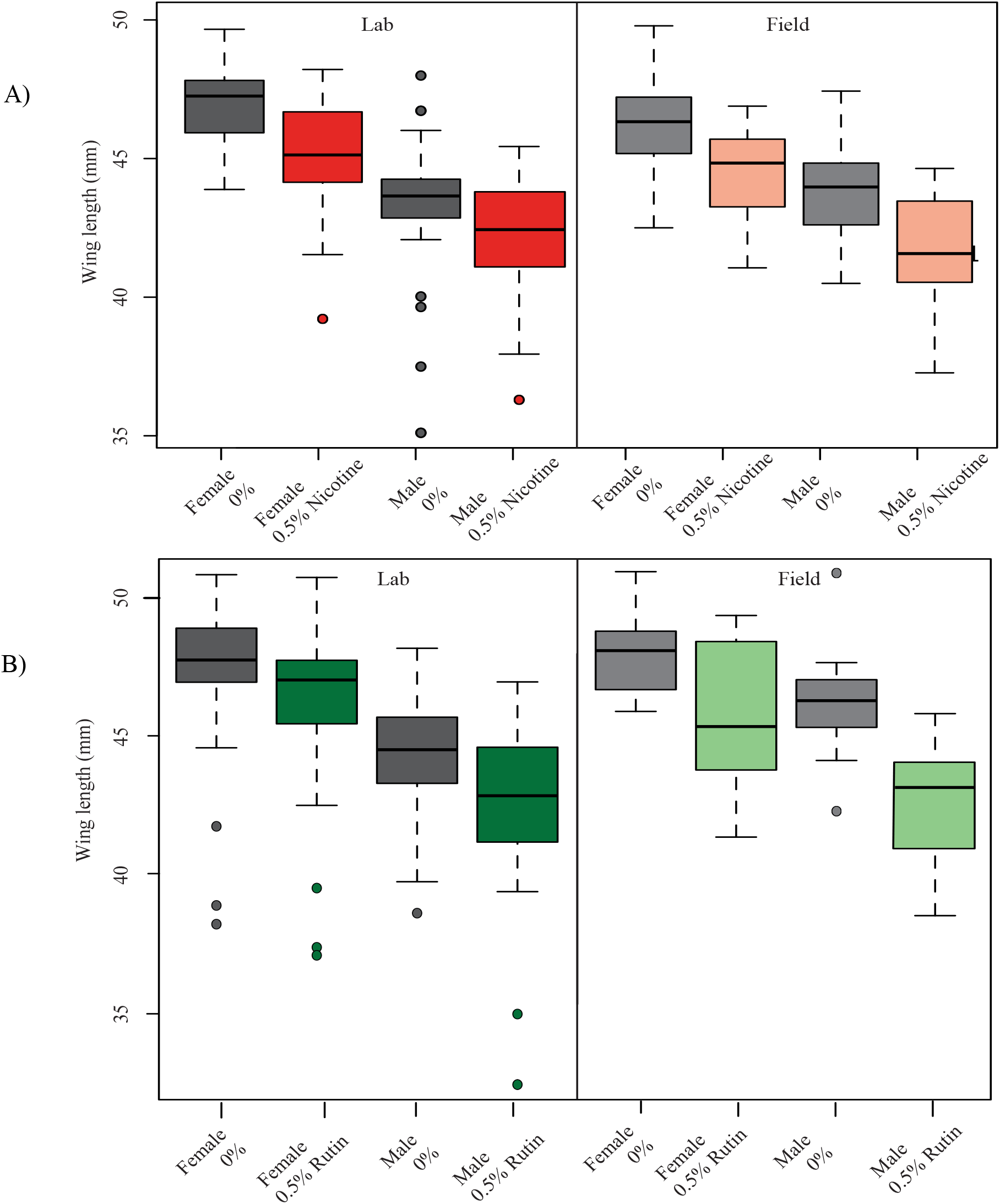
*Manduca sexta* adults had shorter wings when reared on larval diets containing secondary compounds. Males had shorter wings than females but there were no sex-specific effects of either compound (ANOVA *P* > 0.1 for all sex-diet interactions). **A)** Nicotine (red) decreased wing length compared to control diets (grey) in both the lab (left pane, ANOVA *P* < 0.01) and field-collected colonies (right pane, *P*< 0.01). **B)** Rutin (green) decreased wing length compared to control diets (grey) in both the lab (left pane, *P* < 0.01) and field-collected colonies (right pane, *P* < 0.01). Separate controls are presented for nicotine and rutin because the effects of these secondary compounds were tested at different generations.

Similarly, larval rutin consumption decreased adult wing size (ANOVA; field rutin *F*_1, 42_= 20.99, P<0.001; lab rutin *F*_1, 106_ = 7.528, *P* = 0.007). Males had smaller wings than females (field sex *F*_1, 42_ = 15.504, *P*<0.001; lab sex *F*_1, 106_ = 44.040, *P*<0.001) but the effect of rutin did not differ between the sexes within a colony (field sex*rutin *F*_1, 42_ = 1.301, *P* = 0.26; lab sex*rutin *F*_1, 106_ = 0.02, *P* = 0.88) (Fig. 3B).

### Female fecundity was unaffected by larval consumption of secondary compounds

Female fecundity (follicle number) did not differ between *M. sexta* reared on control diets versus those reared on diets containing secondary compounds, even when taking adult size differences into account. Although pupal weight and adult size traits were positively correlated, correlations between adult body size and follicle numbers were not present in most treatments (SI Table 3). Larval consumption of nicotine did not reduce adult follicle-body area ratios (ANOVA; field nicotine *F*_1,39_ = 0.113, *P* = 0.738; lab nicotine *F*_1, 57_ = 0.663, *P* = 0.419). Similarly, larval consumption of rutin did not decrease the follicle-body area ratio (ANOVA; field rutin *F*_1,13_ = 0.085, *P* = 0.775; lab rutin *F*_1,52_ = 0.817, *P* = 0.370).

### Neonate *M. sexta* do not show behavioral avoidance of diets with secondary compounds

In the behavioral experiments testing for a preference for diets with or without secondary compounds, neonate larvae did not display significant avoidance of either nicotine or rutin diets. For larvae that chose between nicotine or control diets (N = 25/30 field, N= 36/38 lab), 48% of the field-collected colony chose control diet (*X*^2^_1_ = 0.04, *P* = 0.842) and 64% of the lab colony chose control diet (*X*^2^_1_ = 2.778, *P* = 0.096). For larvae that chose between rutin or control diets (N = 24/30 field, N = 37/38 lab), 38% of the field-collected colony chose control diet (*X*^2^_1_ = 1.5, *P* = 0.221) and 43% of the lab colony chose control diet (*X*^2^_1_ = 0.676, *P* = 0.411).

## Discussion

Plant-insect coevolution depends not only on plant defenses and herbivore counter-adaptions to these defenses, but also on tri-trophic interactions that include top-down effects (Bruce 2014). In this study, I examined the effects of plant secondary compounds on herbivore fitness in the context of natural enemies. Using field surveys of parasitoid prevalence paired with experimental measurements of the effects of larval consumption of secondary compounds on growth and fitness traits across *Manduca sexta* life stages, I show that natural enemies kill a large proportion of *M. sexta* larvae in the field, while the fitness effects of dietary secondary compound ingestion are less severe (Table 4). Previous studies have established that a defended diet is protective against parasitoids (Thorpe and Barbosa 1986, Barbosa *et al*. 1991, Harvey *et al*. 2007) but the ecological importance of this benefit is highly dependent on parasitoid prevalence. Prior studies using controlled parasitoid oviposition on predetermined hosts or studies that introduce laboratory *M. sexta* into a field setting may not reflect interactions in the field. In this study, I provide important evidence that that parasitoids kill a large proportion of *M. sexta* larvae in the field, representing a total loss of fitness for parasitized hosts.

**Table 4.** Summary of *Manduca sexta* responses to secondary compounds for the traits measured across life stages. Dashes indicate no significant effect and an arrow indicates a significant negative effect of the secondary compound compared to the control diet at *P* < 0.05. The direction of the arrow indicates whether the secondary compound increased or decreased the trait.

| Stage | Trait | effect of nicotine |  | effect of rutin |  |
| --- | --- | --- | --- | --- | --- |
|  |  | field | lab | field | lab |
| larvae | immunity | - | - | - | - |
|  | days per instar | ↑ | ↑ | ↑ | ↑ |
|  | incomplete molting | - | - | - | - |
|  | diet choice | - | - | - | - |
| pupa | development time | ↑ | ↑ | ↑ | ↑ |
|  | pupal mass | ↓ | ↓ | ↓ | ↓ |
| adult | survival | - | - | - | - |
|  | body length | ↓ | ↓ | ↓ | - |
|  | wing length | ↓ | ↓ | ↓ | ↓ |
|  | follicle number | - | - | - | - |

The *M. sexta* larval growth patterns seen in this study minimize exposure to parasitoids during the timeframe that larvae are most likely to be parasitized. I found relatively faster *M. sexta* development time during the second and third instars, which aligns with the instars preferred for *C. congregata* oviposition (Gilmore 1938, Beckage and Riddiford 1978, Barbosa *et al*. 1991, Kingsolver *et al*. 2012). In the field survey, similar numbers of parasitoids emerged from larvae removed from the field as second to fourth instars, suggesting that larvae remaining in the field as fourth instars did not result in additional parasitoids. Parasitoids also emerged quickly from larvae removed from the field as fourth instars, indicating the parasitoids had been laid prior to the fourth instar based on a 12-16 day oviposition-to-emergence time (Gilmore 1938).

Rapid development that reduces exposure time to parasitoids is expected to be beneficial, as fast growth did not come at the cost of reducing immune responses to an artificial parasitoid egg. Although *C. congregata* often lay more than a single egg in an oviposition event, the immune stress represented by a single parasitoid egg represents a parasitoid attack that an *M. sexta* larvae could survive by mounting a strong immune response that prevented the parasitoid egg from hatching. In contrast to the predicted energetic trade-off between growth and immunity (Smilanich *et al*. 2009), I found that larvae with higher growth rates following injection of the artificial parasitoid egg actually had higher levels of melanization regardless of control or defended diets. The lack of a growth-immune trade-off in *Manduca sexta* has also been observed in other recent studies (Wilson *et al*. 2019) and may be because *M. sexta* larvae do not actively sequester plant compounds and therefore do not have this energetic cost (Wink and Theile 2002, Smilanich *et al*. 2009). Nicotine and rutin did not increase melanization of the artificial parasitoid egg, suggesting that host immune responses to secondary compounds are unlikely to be a significant driver of the reduced parasitoid success on hosts fed nicotine seen in other studies (Beckage and Riddiford 1978, Barbosa *et al*. 1986, Barbosa *et al*. 1991, Harvey *et al*. 2007). Although it is possible that real and/or additional parasitoid eggs would increase *M. sexta’s* immune response to a greater extent than an artificial parasitoid egg, *C. congregata* parasitoids have been shown to impair host encapsulation responses by infecting their hosts with immunosuppressant viruses during oviposition (Amaya et al. 2005). Therefore, the protective effects of secondary compounds against *M. sexta* parasitoids probably result from toxicity of nicotine to the parasitoids or the indirect effects of slowed *M. sexta* growth on *C. congregata* development (Barbosa *et al*. 1986, Appel and Martin 1992). Interestingly, I observed higher melanization rates in the field-collected colony than in the lab colony, which may reflect that the lab colony has been removed from parasitoid pressures for many generations (Diamond and Kingsolver 2011, Kingsolver *et al*. 2020).

Secondary compounds had negative effects on larval size and developmental time that can impact interactions with parasitoids, even in the absence of effects on immune responses. Different larval instars may be more or less susceptible to the effects of secondary compounds (van Dam *et al*. 2011) because of the differences in size and amount of food required to complete the different instars. In this study, nicotine and rutin extended the amount of time larvae needed to complete each instar, with no instar specific effects of either compound. Delayed development time and smaller size of *M. sexta* fed secondary compounds as larvae may be the result of changes in digestion, energy spent on maintenance metabolism (Appel and Martin 1992), reduced consumption of defended diets (Voelckel 2001), or disruptions in juvenile hormone (Lee *et al*. 2015). The low levels of incomplete molting I observed in *M. sexta* larvae indicate any changes in hormone levels due to secondary compounds were not enough to fully disrupt molting. Regardless of mechanism, an extended development time means herbivores are exposed to parasitoids for longer when feeding on defended tissues, but this may be offset by the smaller size of these larvae making them harder for parasitoids to locate (Clancy and Price 1987, Benrey and Denno 1997).

These effects of exposure to secondary compounds at the larval stage contribute to *M. sexta* fitness either by altering the probability of surviving to reproduce as adults or through correlations with adult reproductive traits. In the absence of parasitoid pressures, larval consumption of secondary compounds did not affect survival to adult eclosion or fecundity but had negative effects on adult body size and wing size. Females with smaller bodies have been shown to have reduced pheromone production in other Lepidoptera and may be less attractive to males (Harari *et al*. 2011). Wing size is positively correlated with increased flight time and distance which may be important for finding mates or host plants for egg laying (Shirai 1993, Berwaerts *et al*. 2002, Cahenzli *et al*. 2015). Although I found that fecundity (female follicle numbers) did not differ based on consumption of secondary compounds, actual fertility may be lower if these eggs are not fertilized because of reduced mating success. Body and wing traits may also impact an adult’s ability to disperse offspring and/or choose appropriate oviposition sites. However, the impact of maternal oviposition choice depends on whether offspring can behaviorally select their own feeding sites or if early diet is determined by hatching location (Jaenike 1978, Soler *et al*. 2012).

The lack of neonate differentiation I observed between defended and non-defended diets is evolutionarily important because it indicates that maternal oviposition choices rather than offspring choices likely determine whether offspring experience early exposure to secondary compounds (Kester *et al*. 2002). Behavioral responses to plant compounds may vary based on herbivore age. In other studies that have used older larvae, nicotine has shown to be deterrent (Kester *et al*. 2002; Parr and Thurston 1972) while rutin has not been shown to deter feeding and has even been seen to stimulate feeding (De Boer and Hanson 1987, Stamp and Skrobola 1993). The lack of an effect of secondary compounds on neonate choice could indicate that mechanisms needed to recognize chemical cues are not completely developed until later instars, that there are additional important leaf cues that are not present in an artificial diet, or that deterrence may occur via post-ingestive mechanisms rather than pre-ingestive mechanisms (Glendinning 2002).

## Conclusions

Overall, the results of this study indicate that although nicotine and rutin differ in their chemical composition and prevalence in plants, both have negative effects on *M. sexta* that extend past the larval stage at which the compounds are consumed. Despite effects on adult body and wing size that may influence mating and offspring dispersal, there were no strong effects on survival or fecundity. Therefore, the negative effects of secondary compounds on *M. sexta* development and fitness are likely outweighed by the known protection that ingestion of these compounds offers against *C. congregata* parasitoids, which exerted large negative survival costs on *M. sexta* in the field. At a larger scale, coevolutionary and tri-trophic interactions can maintain a balance between the costs and benefits of secondary compounds on herbivore fitness.

## Notes

### Competing Interest Statement

The authors have declared no competing interest.

