## Supplementary material for "*Manduca sexta* experience high parasitoid pressures in the field but minor fitness costs of consuming plant secondary compounds": SI Table 1

**SI Table 1.** Recipe for artificial diet used for *Manduca sexta* larval rearing modified from the University of Madison Wisconsin Department of Entomology diet. Experimental and control diets were refrigerated and kept for no longer than one week to prevent degradation of the chemicals over time. For the growth trials, artificial diet was refreshed or replaced every 1-2 days. For the choice trials using neonate larvae, fresh diet was made the day of the trial. To create 0.5% rutin and 0.5% nicotine diets, pure liquid nicotine or powdered rutin were added to diets once cooled but before hardening. The amount of water for control diet (530 mL) was adjusted for the experimental diet to maintain consistency with the addition of 0.5% nicotine (liquid) or 0.5% rutin (powdered). Artificial diet was blended with 10-20 percent fresh leaf tissue for the larvae collected in the field (tobacco in 2013 and *Datura stramonium* in 2014) as larvae will not consume artificial diet lacking plant material after feeding on leaves.

| **Ingredient** | **Amount** |
| --- | --- |
| Water | 530 mL* |
| Agar | 20 g |
| Non-toasted wheat germ | 100 g |
| Non-fat dry milk | 22 g |
| Sugar | 13.6 g |
| Vitamin C (generic) | 1 tablet |
| Vitamin B (generic) | 2 tablets |
| Multivitamin (generic) | 2 tablets |
| Raw flax seed oil | 1 tsp |
| Nutritional flake yeast | 3.8 g |
| Sorbic acid (preservative) | 1.5 g |
| Methyl-4-hydrobenzoate (preservative) | 1 g |
