## Supplementary material for "*Manduca sexta* experience high parasitoid pressures in the field but minor fitness costs of consuming plant secondary compounds": SI Table 2

**SI Table 2.** Secondary compounds increased the amount of *Manduca sexta* larvae completing an extra (sixth) instar (N = 80 per treatment and colony). Numbers of larvae completing six instars are presented for control and nicotine fed larvae (left) and control and rutin (right) for the field and lab colonies. Significant *P* values (< 0.05) from Fisher’s exact tests are indicated with an asterisk (*).

|  | **control** | **nicotine** | **P** | **control** | **rutin** | **P** |
| --- | --- | --- | --- | --- | --- | --- |
| **Field** | 5 | 2 | 0.44 | 0 | 6 | 0.03* |
| **Lab** | 0 | 6 | 0.02* | 0 | 8 | <0.01* |
