## Supplementary material for "*Manduca sexta* experience high parasitoid pressures in the field but minor fitness costs of consuming plant secondary compounds": SI Methods 1

**Supplementary methods 1:** Establishment of a field-collected colony

Once surviving field-collected *M. sexta* reached the pupal stage, they were transferred to a large cage to eclose and mate in order to establish a field-collected colony to use for the experiments testing the effects of secondary compounds on *M. sexta* developmental and fitness. Pupae were placed over a tray of wet gravel in a greenhouse with 14D:10N light cycles and misted daily with water to increase humidity in the cage. The cage was draped with a sheet to keep the moths cool and increase darkness at night. Adult moths were provided with a sponge saturated with 20% honey water for nectaring (Hunter’s honey farm, Martinsville IN, USA) and a Solanaceous host plant for egg deposition. Bergamot oil was also applied to the cage every 1-3 days to increase nectaring and stimulate mating (Goyret and Raguso 2006). Eggs produced from the matings were collected and surface sterilized with 1% bleach and rinsed in distilled water. After hatching, *M. sexta* larvae were reared individually in two-ounce clear plastic lidded containers until approximately the third instar, when they were transferred to similar lidded four-ounce containers.

Although *M. sexta* larvae directly collected from the field required the addition of leaf tissue to the artificial diet, larvae of subsequent generations in the lab were reared solely on artificial diet (SI Table 1). Approximately 60 percent of the larvae take to the artificial diet while the remaining die within the first two to four days after hatching after failing to accept the artificial diet.
