## Supplementary material for "*Manduca sexta* experience high parasitoid pressures in the field but minor fitness costs of consuming plant secondary compounds": SI Figure 1

**SI Figure 1**. Fate of *Manduca sexta* larvae collected from a cultivated field of *Nicotiana tabacum* in Kentucky, USA at three time points. A total of 395 *M. sexta* larvae were collected from the field over the three collection dates (21 July 2013 N=98; 20 August 2013 N=156; 28 July 2014 N=141). The parasitoids emerging from field-collected *M. sexta* were *Cotesia congregata* (“larval parasitoids,” black band) and Tachinid flies (“pupal parasitoids,” small dark gray band present in July 2013 and July 2014). *Manduca sexta* larvae that did not survive but did not show external evidence of parasitoids are indicated by the light gray band. The larvae that survived and pupated as adults in the lab (white band) were mated and used to generate the experimental field-collected colony in the greenhouse.

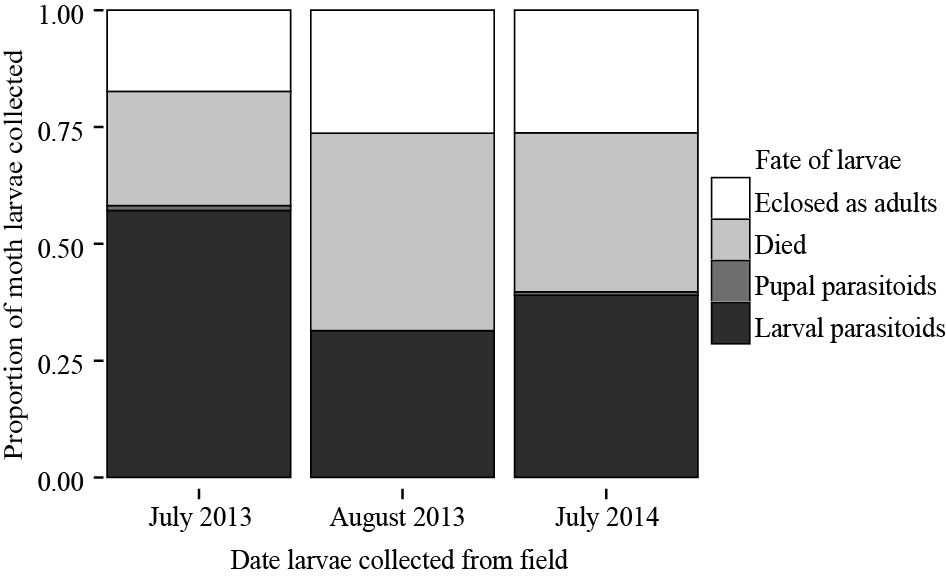
